# Control limitations in the null-space of the wrist muscle system

**DOI:** 10.1101/2023.11.28.568721

**Authors:** Meng-Jung Lee, Jonathan Eden, Sergio Gurgone, Denise J Berger, Daniele Borzelli, Andrea d’Avella, Carsten Mehring, Etienne Burdet

## Abstract

The redundancy present within the musculoskeletal system may offer a non-invasive source of signals for movement augmentation, where the muscle-to-force null-space could be controlled simultaneously to the natural limbs. Here, we investigated the viability of extracting movement augmentation control signals from the muscles of the wrist complex. Our study assessed i) if controlled variation of the muscle activation patterns in the wrist joint’s null-space is possible; and ii) whether force and null-space targets could be reached simultaneously. During the null-space target reaching condition, participants used EMG-to-force null-space muscle activation to move their cursor towards a displayed target while minimising the exerted force as visualised through the cursor’s size. Initial targets were positioned to require natural co-contraction in the null-space and if participants showed a consistent ability to reach for their current target, they would rotate 5° incrementally to generate muscle activation patterns further away from their natural co-contraction. In contrast, during the concurrent target reaching condition participants were required to match a target position and size, where their cursor position was instead controlled by their exerted flexion-extension and radial-ulnar deviation, while its size was changed by their natural co-contraction magnitude. The results collected from 10 participants suggest that while there was variation in each participant’s co-contraction behaviour, most did not possess the ability to control this variation for muscle-to-force null-space reaching. In contrast, participants did show a direction and target size dependent ability to vary isometric force and co-contraction activity concurrently. Our results show the limitations of using null-space activity in joints with a low level of redundancy.

## Introduction

Movement augmentation technology promises to increase a user’s effective mechanical degrees-of-freedom (DoFs). For instance, a surgeon could use a supernumerary robotic limb to perform laparoscopy without an assistant^1^, or a user-interface could enable a person to control their phone while their hands carry heavy luggage. However, an increase in DoFs comes with a corresponding need to control them. For practical DoF augmentation to be achieved it is therefore necessary that i) the user is able to control the extra DoFs with a functional level of accuracy; and ii) this control has minimal interference on existing natural motion behaviour. While movement augmentation has been demonstrated for a number of different applications^2^, the limits and mechanisms for its practical application are still unclear.

A number of different control schemes have been developed to support human DoF augmentation^3^. These typically make use of autonomous augmentation^4,5^, in which robot DoFs are controlled without the user’s volitional control, or augmentation by transfer^6,7^, where body DoFs not relevant to the task are substituted to control the extra DoFs. While such approaches can enable task specific DoF augmentation, their application is limited. Here, autonomous augmentation requires that the robot can predict the operator’s desired actions (which can lead to poor functional accuracy with respect to the user’s intent). Meanwhile, augmentation by transfer impacts natural motion unless it is restricted to use cases that do not use the substituted body DoFs.

Augmentation by extension, which uses body signals that are not involved in motion, could enable DoF augmentation that is application independent^3^. While invasive methods for simultaneous brain computer interfaces (BCI) and natural movement control have been investigated^8,9^ and could potentially be used as augmentation by extension schemes, non-invasive approaches (e.g, electroencephalogram^10^ or motor unit recording from high-density surface electromyography (EMG)^11–13^) can be more accessible and safer.

An alternative possible non-invasive augmentation by extension approach is to use the redundancy naturally present within the musculoskeletal system through the task-intrinsic muscular null-space^14^. The human body possesses more muscles than effective mechanical DoFs such that there is redundancy if those additional muscles can be controlled in atypical patterns to produce the same task-space force. This redundancy could potentially provide natural and measurable signals for human movement augmentation. Here, it has been shown that after remapping muscle activation to new muscle synergies, humans can (with sufficient exploration time) adapt their muscle activation behaviour in response to either simple remappings requiring combinations of existing muscle synergies, or the more challenging case requiring completely new synergies^15^. Moreover, it has been found that muscle activation patterns that fall into the muscle-to-force ‘null-space’ during isometric movement can be used as an additional control signal that is regulated simultaneously to the generation of isometric force^16^. These results suggest the potential for using muscle redundancy as a source of command signals for extra DoFs. However, while it has been shown that simultaneous null-space and reaching actions can be performed when the redundancy is from the entire limb to end-effector forces^16^, it is difficult to evaluate if natural body behaviour has been altered due to the complex non-linear biomechanics involved and it is unclear if these results are only possible with the large levels of redundancy present across the entire upper limb.

In this study we explored if the redundancy from the muscles of the wrist joint to its 2 revolute DoFs could be used to extract control signals for DoF augmentation. Using EMG signals recorded from 4 wrist muscles (ECRL, ECU, FCR, FCU) we conducted an experiment with 2 sessions considering if: i) humans can alter their natural co-contraction behaviour to produce atypical muscle activation patterns that do not produce force; and ii) they can simultaneously vary both task-oriented reaching motions and activity that is in the null-space of their force production. This used an incremental training paradigm to optimize the participants’ learning process^17^.

## Methods

### Participants

The experiment was approved by the Research Governance and Integrity Team at Imperial College London (Reference: 21IC6935) and was carried out by 10 participants (5 female, 5 male) without known sensorimotor impairment aged 24-36. All participants partook in both experiment sessions. 1 participant’s data from the concurrent target reaching session was removed due to a computer crash during the session. Each participant had no prior experience with the experimental paradigm and gave their informed consent before participating. All participants self-reported to be right-handed and had normal or corrected to normal vision.

### Experimental setup

The experiment was conducted using a fixed handle (Fig. 1A,^18^) attached to a 6 DoF force torque sensor (Nano SI-25, ATI Industrial Automation). The seated participants had their right arm fixed to the interface at the wrist, forearm and elbow. Surface electromyography (EMG) was recorded using the g.GAMMASYS system (g.tec) from 4 wrist muscles: flexor carpi radialis (FCR); flexor carpi ulnaris (FCU); extensor carpi radialis longus (ECRL); extensor carpi ulnaris (ECU) and interossei muscles. These 4 muscles are primarily responsible for 2 mechanical DoFs (flexion-extension and radial-ulnar deviation). Mechanically, to produce any single force in these 2 dimensions, at most only 2 muscles are required to be active. As a result for any produced force, the wrist system possesses 2 potential degrees of redundancy resulting from the activation of its other 2 muscles. The recorded muscles were located manually at the suggested sites described in^19^ together with palpation. The interface and EMG recording was operated at 1000 Hz, where the EMG signals were first high pass filtered with a cut-off frequency of 20Hz, then rectified and finally low-pass filtered with a 5Hz cut-off frequency (all second-order Butterworth filters).

**Figure 1.**
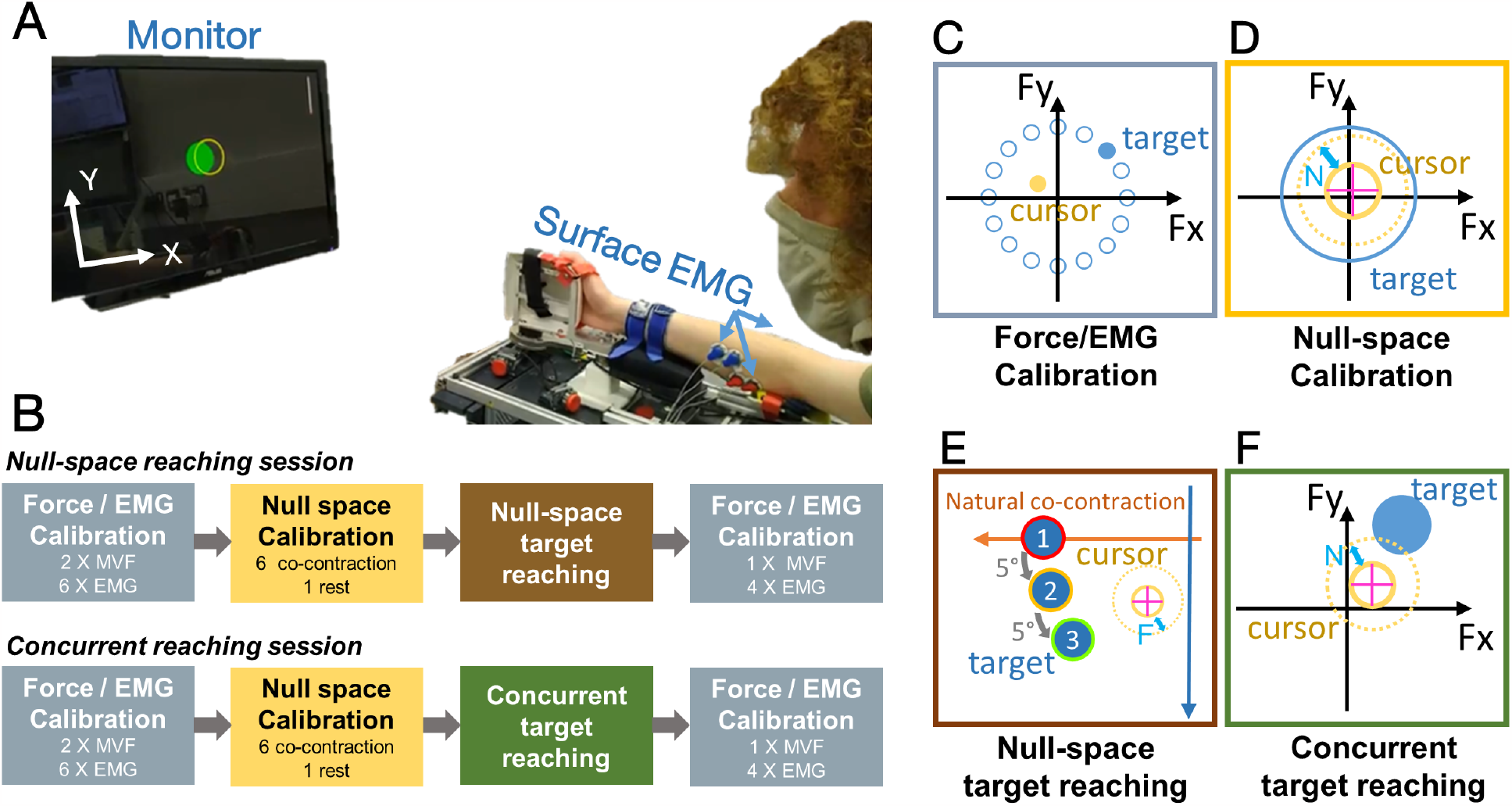
Experiment setup and design. (A) Participants were attached to an ergonomic handle mounted to a force torque sensor and were required to reach for targets using the recorded force signal and/or surface EMG recordings. (B) The experimental protocol for the 2 experiment sessions, where participants completed both sessions in a randomised order. A visualisation for each block is shown in (C)-(F). Depending on the block, participants controlled the position (indicated by a purple cross on the yellow cursor) and size (D)-(F) of the cursor to match different targets (shown in blue). In the NSTR (E), the cursor position was controlled by activation in the EMG-to-force null-space where natural co-contraction was mapped to negative X-axis motion, in all other blocks force control was used. Possible target locations are shown in the Force/EMG calibration (C) and null-space target reaching blocks (E).

### Experiment design

To investigate the ability of participants to exploit the redundancy present within their wrist system, the vector of 4 recorded EMG signals was (at each time step across the experiment) projected onto a 2-dimensional *task-space* component that resulted in the production of flexion-extension and radial-ulnar deviation force (computed using the method of^20^) and a 2-dimensional *null-space* component that did not result in force/torque (as in^16^). The experiment was conducted across 2 different sessions: *null-space target reaching* (NSTR) and *concurrent target reaching* (CTR), which are defined below. During the NSTR session, the participants’ ability to control only the null-space component of their EMG activation pattern without producing force was evaluated. Here, participants were required to produce a null-space activation of a fixed magnitude in different locations of the null-space, which would require them to alter their EMG activation patterns so as to deviate from their natural co-contraction behaviour without producing force. In contrast in the CTR session, the participants ability to produce null-space activity concurrently to force/torque activation was evaluated by requiring participants to reach for targets in different locations of the task-space while generating the correct amount of activation in the null-space through co-contraction.

The experiment took place over 2 days with no more than 1 day of break in between. Participants completed 1 session each day and were randomly (with equal probability) assigned to start with either the NSTR or the CTR session. Fig. 1B depicts the experiment protocol for each session. Here, each (approximately 90 minute session) consisted of the same series of experimental blocks: an initial *force/EMG calibration*, a *null-space calibration*, a main experiment block and a final post session force/EMG calibration to confirm that there was no changes induced by the main block. The sessions therefore only differed in the different main experiment blocks.

### Tasks

#### Force/EMG calibration

The participant specific *maximum voluntary force* (MVF) and the mapping from EMG activation to task-space force (or *pulling vector* matrix) were computed. Throughout this phase, the participants were instructed to face the monitor which displayed a yellow cursor representing the measured *X* (extension-flexion) and *Y* (radial-ulnar deviation) axis force. They were then instructed to one-at-a-time provide a maximum exertion of force in 1 of 16 equally spaced (every 22.5°) directions (Fig. 1C), where the reference direction was displayed on the monitor with a dashed line and progressed anticlockwise starting initially from leftward (flexion) force production. Each direction was given 4 seconds of time and the MVF for each direction was computed as the maximum recorded force from the last 2 seconds, and the participant specific scaling MVF (sMVF) was set to be the minimum recorded MVF value across the 16 directions. The force visualisation was then scaled by the sMVF such that this magnitude of force would move the cursor to the boundaries of the monitor. With this scaling the pulling vector matrix was calibrated for using a similar procedure to^20^. Participants were asked to produce and hold (for 2 s) their force at 17 different target locations, corresponding to 1 target at rest with [0, 0] N force and the remaining 16 targets evenly spaced along a circle with radius 0.2 sMVF. For each target, the average of the force **f** = [*f*_*x*_, *f*_*y*_]^*T*^ and EMG activity ***α*** = [*α*_*FCR*_, *α*_*FCU*_, *α*_*ECRL*_, *α*_*ECU*_]^*T*^ was recorded across the 2 s holding period. After multiple rounds of targets, the pulling matrix **H** *∈* ℝ^2×4^ was computed in a manner consistent with^21^ through ordinary least squares (see Supplementary Figure S1 for a visualisation of the pulling vector matrix) assuming the linear relationship

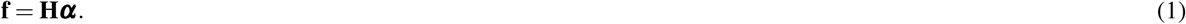

The initial calibration and post session calibration blocks contained an identical process, however, to account for participants not having familiarity with the calibration procedure, the initial calibration was composed of 2 rounds of MVF production and 6 rounds of EMG calibration target reaches, while the post calibration consisted of only 1 round of MVF production and 4 rounds of target reaches.

### Null-space calibration

The null-space with respect to the produced force was computed and the components of its visualisation were oriented. First the null-space projection matrix **N** was determined such that its rows consisted of orthonormal basis vectors that were orthogonal to the subspace projected by the pulling matrix **H**. Then, a scaling

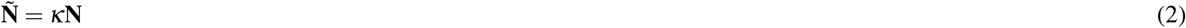

was applied to ensure that the resulting null-space coordinates

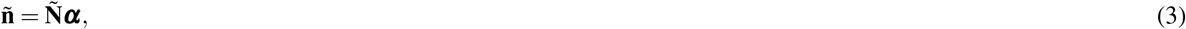

had a similar magnitude to that of the produced force (for the same muscle activity levels), where **ñ** = 1 would have similar EMG activity magnitude to that of 1 N of produced force. This scaling was determined so that its magnitude was equal to the largest of the scaling from EMG activity to the principle force directions (flexion-extension or radial-ulnar deviation), such that *κ* = max {∥ *h*_1_∥, ∥ *h*_2_ ∥}, where *h*_*i*_ denotes the ith row of the pulling matrix **H**. With a null-space mapping computed, the cursor was augmented such that in addition to controlling its position through the sMVF scaled *X* and *Y* axis force, participants also were in control of the cursor radius through the null-space component **ñ**’s magnitude.

To determine the natural direction that co-contraction took within the null-space, participants were then instructed to “naturally” stiffen their wrist such that the cursor remained at the origin position while its size grew to match a target ring with ∥ **ñ** ∥ = 3 (Fig. 1D). Participants were required to repeat this task 6 times, where for each repetition the last 2 s of EMG activity was recorded (see Supplementary Figure S2 for a visualisation of the average EMG activity in each of the 6 repetitions) and used to compute the average co-contraction null-space activity **n**_**c**_. Participants were then asked to perform 1 final calibration trial in which they were instructed to completely rest. This was used to determine their offset null-space activity **n**_**0**_. From the null-space offset and the average co-contraction null-space activity **n**_**c**_, the null-space component **n** was transformed such that it was **0** at rest and natural co-contraction corresponded to horizontal motion on the screen. This meant that the final null-space activity **n** was given by

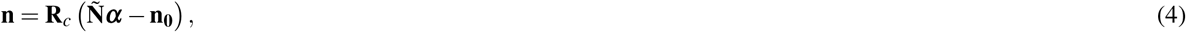

where **R**_*c*_ corresponds to the matrix that rotates the vector **n**_**c**_ to [1, 0]^*T*^.

### Null-space target reaching

The visualisation was changed such that the *X −Y* position on the monitor corresponded to the negative of the null-space activity **ñ** with natural co-activation leading to leftward motion, and the radius of the cursor was proportional to the magnitude of the recorded force. Before the participants were asked to attempt 80 null-space target reaches, they were first given 2 minutes of exploration time to freely explore and get familiar with the altered visualisation without specific instruction. For each of the 80 targets, the participant had 7 s to reach for the target location in the null-space. A target reach was considered successful if the participant reached its location and held at that location for 1 s without leaving the target or producing more than a predefined force threshold (0.04 sMVF). The cursor was considered in the target if the null-space error **e**_*ñ*_ between the target and cursor location was less than a predefined threshold (*∥***e**_*ñ*_ *∥≤* 0.4). If the cursor satisfied this condition while being below the target threshold, the target colour would change to green to indicate that both conditions were met.

The target was initially located to correspond to the natural co-contraction measured during null-space calibration. Therefore it was placed at [2, 0]^*T*^ (which due to the flipped coordinates was visualised on the left side of the origin). Each successive target reach would use the same location until the participant demonstrated that they could consistently reach the target, which was defined as successfully reaching at least 80% of targets over a moving window of ten trials. If this condition was met, a new target location would be triggered corresponding to an anticlockwise rotation of the previous location by 5° (Fig. 1E). This was set such that each new target would require increasing the null-space activity in the direction that was orthogonal to natural co-contraction, while being small enough so that participants needed small adaptation for successive targets. At best the participants could reach for targets in ten different target locations within the 80 reaching trials.

### Concurrent target reaching

The visualisation was the same as in the null-space calibration. Participants were required to make 90 target reaches from the centre to the target, where as in the NSTR, they had 7 s to reach for the target location and would be considered successful if they held at that location for 1 s. In this task, targets were defined both in terms of their task-space location (visualised as the target’s position on the screen) and the natural co-contraction coordinate of the null-space (visualised as the target’s size). Fig. 1F illustrates the visual feedback for the participants. 9 different task-space locations were considered corresponding to the origin and 8 equally spaced points on a circle with magnitude 0.2 sMVF, while 2 different sizes were considered corresponding to null-space activity∥ **ñ** ∥ = 3 and ∥ **ñ** ∥ = 6. Target sizes and locations were randomised and could change for each target reach. If the cursor was within the tolerances in reaching the target, the target changed colour to be green. Before each reach, participants were required to return to the centre position to initiate the next trial.

### Data analysis

The participant behaviour was evaluated in terms of the *task performance* and *motion characteristics* for each experimental session. In the NSTR, task performance consisted of the number of different target locations that were attempted (*number of targets*) as well as the participant’s displayed angular range of motion within the null-space (*reaching range*). The reaching range was individually computed for the practice phase and the main experimental phase as well as the union of the data from both phases. In contrast, since the CTR had a fixed number of trials for each target location, task performance was evaluated through the percentage of successful trials (*success rate*) for each target.

The participant’s motion characteristics were evaluated in the NSTR through 4 metrics: i) the *motion efficiency* computed as the ratio of the direct straight line motion to the total distance travelled in the null-space for a successful target reach; ii) the *time to success* defined as the time interval between the start of the trial and the time of a successful reach; iii) the *number of corrections* which was estimated by first low-pass filtering the null-space trajectories (second-order Butterworth; 3 Hz cut off) and then computing the number of peaks in the speed profile (using the MATLAB findpeaks function); and iv) the *force magnitude*, computed as the root mean square of the measured force across the trial.

In the CTR session, the motion characteristics were instead computed through the *time to success* as well as the *motion concurrency* which was computed as the percentage of the total null-space motion that was performed at the same time as the task-space motion. This percentage was computed considering the effect of unintended natural null-space activity which was removed before the subsequent percentage calculation. Here, we used the EMG recording from the equivalent target position reach during the calibration phase (for each individual participant) as the activation baseline (without intentional null-space activation) and predicted how much null-space movement each participant would have generated when reaching for that individual task-space target without any additional co-contraction. The remaining null-space movement from the baseline to the target size was then separated into 2 phases: Phase 1 running from movement onset until the target in task-space was reached; and Phase 2 running from when the task-space target was reached (end of phase 1) to the null-space target reach. The sum of null-space movement from Phase 1 and Phase 2 was equivalent to the difference of the target size and the baseline. If the needed null-space movement was less than 15 % of the target size, it was considered to be too small an effort for any robust measurement and therefore removed from the analysis (8 and 6 data points removed from small and big target size, respectively).

### Statistical analysis

Bayesian statistical techniques were used for hypothesis testing. Outcomes of hypothesis tests are reported by Bayes Factors that quantify the relative support of the data for 2 competing hypothesis. To compare the range of movement angles that participants could generate between their initial null-space exploration and the main null-space target reaching the Bayesian Wilcoxon signed rank test^22^ was applied as a Bayesian alternative of the non-parametric Wilcoxon signed rank sum test. A Bayes factor robustness analysis was performed that demonstrated that the reported Bayes factors remained relatively stable across a range of prior widths. For all other metrics in the NSTR, due to each participant reaching a different amount of target locations such that there are inconsistencies in the amount of data for each target, only descriptive stats are presented.

The Bayesian repeated measures ANOVA^23–25^ was applied to compare the means of multiple dependent samples as a Bayesian variant of the repeated measures ANOVA. This was used to analyse (1) the dependence of the success rate and latency in the CTR on target location and size and (2) the dependence of the activity in the null-space during the CTR on the target size and task phase. In these two-way repeated measures ANOVA designs (with interaction) each effect is included in multiple models. We, therefore, report the inclusion Bayes Factors (*BF*_*inclusion*_) that provides the relative support for all models that include an effect compared to all models excluding it^24,25^. Post-hoc Bayesian paired t-tests were performed to identify pair-wise differences.

All hypothesis tests were performed using JASP 0.17.1 (https://jasp-stats.org).

## Results

### Null-space target reaching

#### Task Performance

Participants showed a mix of abilities for successfully reaching for the null-space targets. Here as shown in Fig. 2A only 6 out of 10 participants were able to show sufficient consistency in their reaching to progress beyond the first target. Of those participants that did progress beyond that target, only 2 were able to progress to attempt the fifth target, while the remaining 4 did not progress beyond the third target. Here it is noted that due to the target size, the first 3 targets could have been successfully reached using the same strategy as the first target. This therefore indicates that the majority of the participants did not show a consistent ability to change their null-space activity.

**Figure 2.**
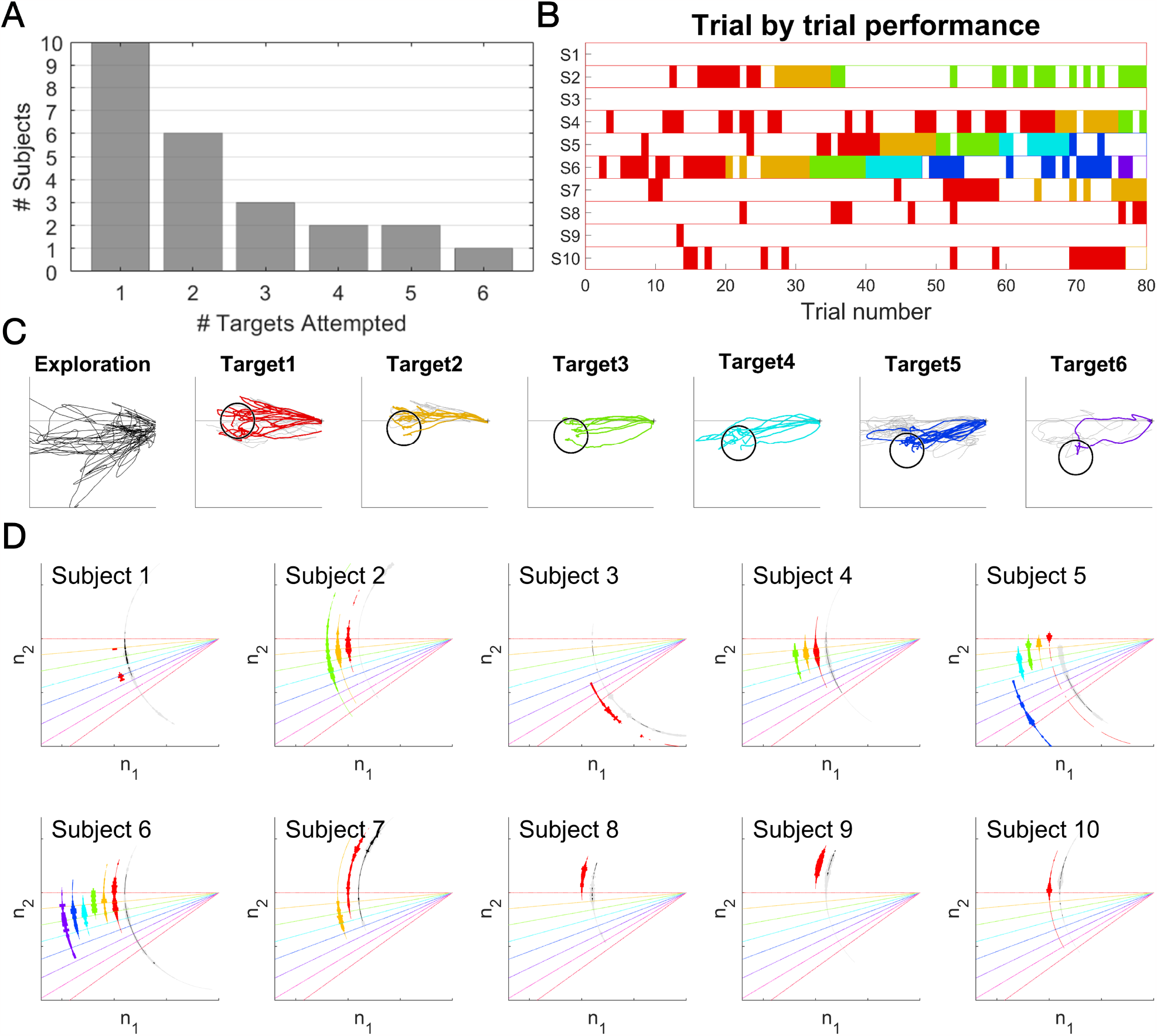
Performance during null-space target reaching. (A) Number of participants that attempted the target location. (B) Distribution of successful trials for each participant. A successful trial is shown by a coloured bar, where each colour represents a different target location (with colours matching those shown in (C)). 6 out of 10 participants met the criteria to move past the first target. (C) Overlaid cursor trajectories from the participant that reached the most targets (Subject 6) during the exploration and target reaching tasks. The black empty circles indicate the target position. Trajectories from failed and successful trials in the target reaching task are shown in grey and coloured lines, respectively. Note that trajectories are only shown up to the time instant at which the target was reached in the successful trial cases. (D) Histogram showing the angles reached for each target location. Angular coordinate show the reached angles from exploration (black: reached angles with the same constraints as in the target reaching task; grey: reached angles that violated the force magnitude constraint) or target reaching task (in colour). Width shows the frequency that the angles were reached. The angle range was larger than the distance between targets, suggesting a potential to move to reach more angles. Colour representation for each target: Red (Target 1); Orange (Target 2); Green (Target 3); Cyan (Target 4); Blue (Target 5); Purple (Target 6).

The overall participant ability is further illustrated through their trial-by-trial success (Fig. 2B). This shows that participants were in general unable to successfully reach for the first target within the initial trials. This was despite that target being set to match their average ‘natural’ co-contraction behaviour across the null-space calibration. Since participants were only able to proceed beyond the second target if they were able to adapt such that they could consistently produce the necessary null-space activation to reach and hold the first target, the use of the number of targets attempted as a metric is limited in that it does not provide understanding of any modulation that may have taken place without successful reaching of the first target (for example if the participant was able to modulate their null-space activity but could not hold it at the required location).

For the best performing participant (Subject 6), Fig. 2C shows their overall null-space trajectories for each target, in which the successful trials are shown coloured and the non-successful trials shown in grey. From these trials it can be observed that this participant showed an ability to move within a large range in the null-space exploration. For the first target, they were able to restrict this motion such that it was mainly horizontal (corresponding to ‘natural’ co-contraction). While similar trajectories were then used for the second and third targets, the participant was able to redirect their motion for the fourth and subsequent targets. However, by the fifth target they showed signs of difficulty for generating further changes in their motion angle. This suggests that in this case they may have reached the limit of their range of controlled motions.

To further investigate the range of null-space activation that the participants were able to generate without the production of force, Fig. 2D shows a histogram of the different angles within the null-space that were reached without violation of the force and null-space magnitude constraints for each participant. Note that the exploration phase component shows both the reached angles with (black) and without (grey) considering the constraint on the force magnitude, respectively. From the exploration phase shown in this figure it can be observed that while participants could often reach a wide range of angles in the null-space, this was typically coupled to the production of force. Furthermore, in the reaching phase participants showed a mixed ability to successfully reach for the different targets. Here, their distribution of motion appears to have a trend that differs for each target that they attempted to reach. However, more data would be needed for each target to validate this trend.

#### Motion Characteristics

The characteristics of the participant motion is further illustrated through Fig. 3. Here, Fig. 3A indicates that participants possessed some range of motion within the null-space exploration. This range of motion appears related to the range of reaching angles observed during target reaching, where there was weak evidence to suggest that there was no difference between the range of angles reached in exploration and in target reaching (*BF*_10_ = 0.38, 0.3, 0.22 for normal, wide and ultra-wide prior with Cauchy prior distribution set to *γ* = 0.5, 1 and 2, Bayesian Wilcoxon signed-rank test).

**Figure 3.**
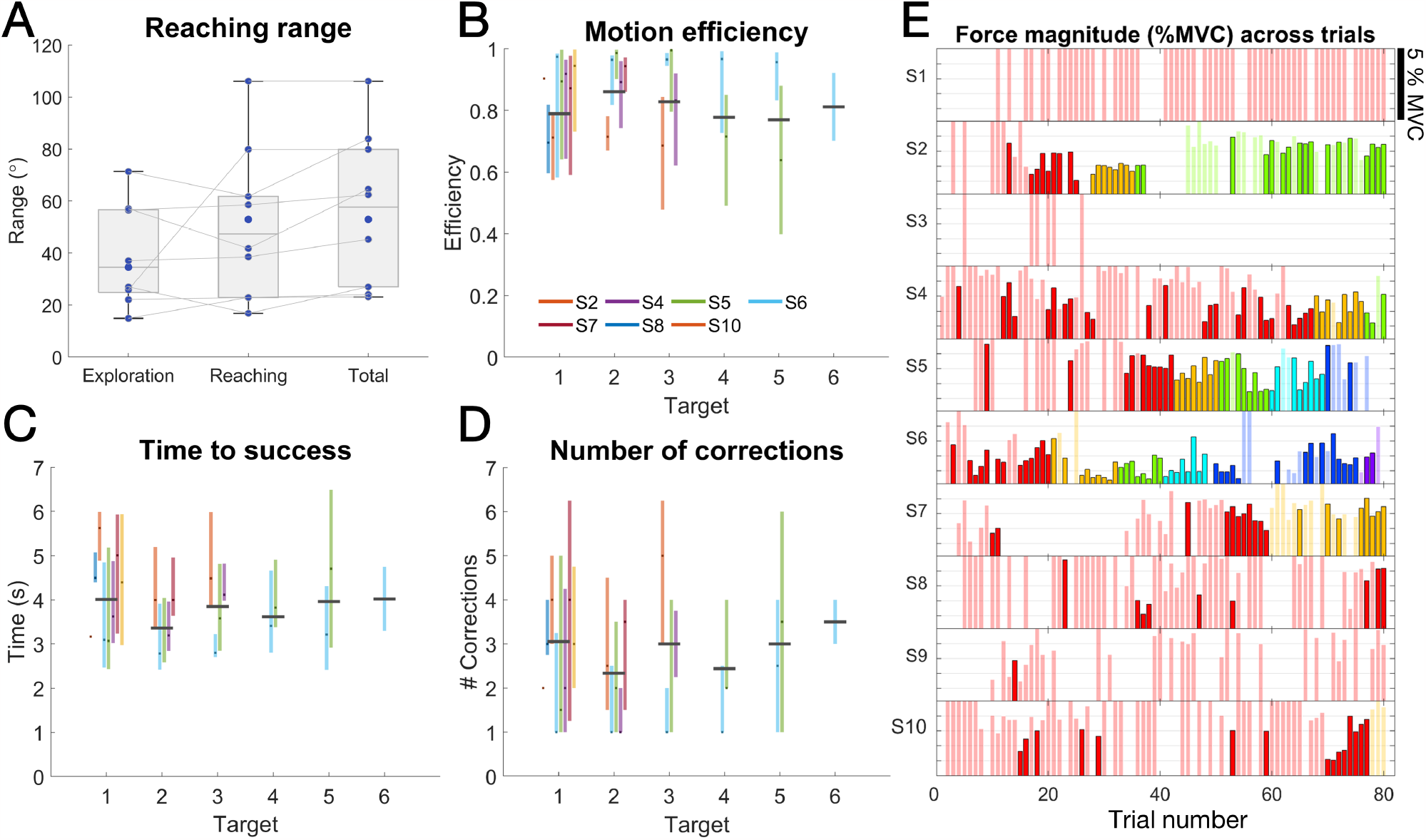
Movement characteristics during NSTR (A) The range of different angles reached for the exploration, null-space reaching phase and across the entire session. (B-D) Motion characteristics of the movement in null-space. Each participant is encoded with a certain colour. Each coloured bar represents the performance of a single participant within the trials of a given target, with the middle being the median and the length being the interquartile ranges. Black horizontal bars over the coloured bars depict the median performance of all participants for the same target. (B) Motion efficiency was computed as the ratio of the Euclidean distance from the cursor’s initial position to the target and the sampled motion’s path length. Most participants show high efficiency suggesting relatively direct motion to the target. (C) The time to success is relatively constant across all the targets. (D) Number of angle corrections from the initial direction (for successful trials this stops when the target is reached) is also relatively constant. (E) Task-space force magnitude when the cursor was at the correct target position for each trial for an individual participant. Saturated colours depict successful trials in the success condition. The transparent colours illustrate the force produced when the cursor is at the target for unsuccessful trials. Each colour is coded for an individual target. Note that the force is only shown until the threshold of 5% MVC. Force magnitude does not increase with trial or target, indicating that the participants were not using higher force level to compensate the null movement.

When considering the successful trials, the motion efficiency (Fig. 3B) shows that in general participant motion was direct towards the target for all locations, where the median efficiency across participants (who reached that target) was greater than 0.75 for all targets. The time to success (Fig. 3C) and the number of corrections made (Fig. 3D) show no clear trend of change across the target locations. However, it should be noted that the small number of participants that reached the targets with larger reaching angles limits this analysis.

Finally, while participants showed some force production in their null-space activation, most participants were able to keep it below the threshold, and within successful trials they do not appear to have an increase in the magnitude of force produced (Fig. 3E) with respect to the different target locations.

### Concurrent target reaching

To test the feasibility of performing null-space target reaching simultaneously with movement in the task-space, participants used their 2-dimensional wrist force to reach for a target at 1 of the 9 locations, with target T9 corresponding to no force needed (Fig. 4A). Concurrently they were required to use their null-space activation projected onto their natural co-contraction to reach the given target size (∥**n**∥= 3 or ∥**n** =6∥), which corresponded to the 2 different levels of null-space activation levels. Only when both target position (task-space) and size (null-space) were reached jointly would the trial be considered as successful. The participant success rate depended on target location and size (Bayesian repeated measures ANOVA, *BF*_*inclusion*_ = 9.9, 3.0, 6.6 for target size, location and interaction of size and location) with higher success rates when reaching for the small targets and lower success rates for the large targets in the ulnar directions (T6-8) (Fig. 4B).

**Figure 4.**
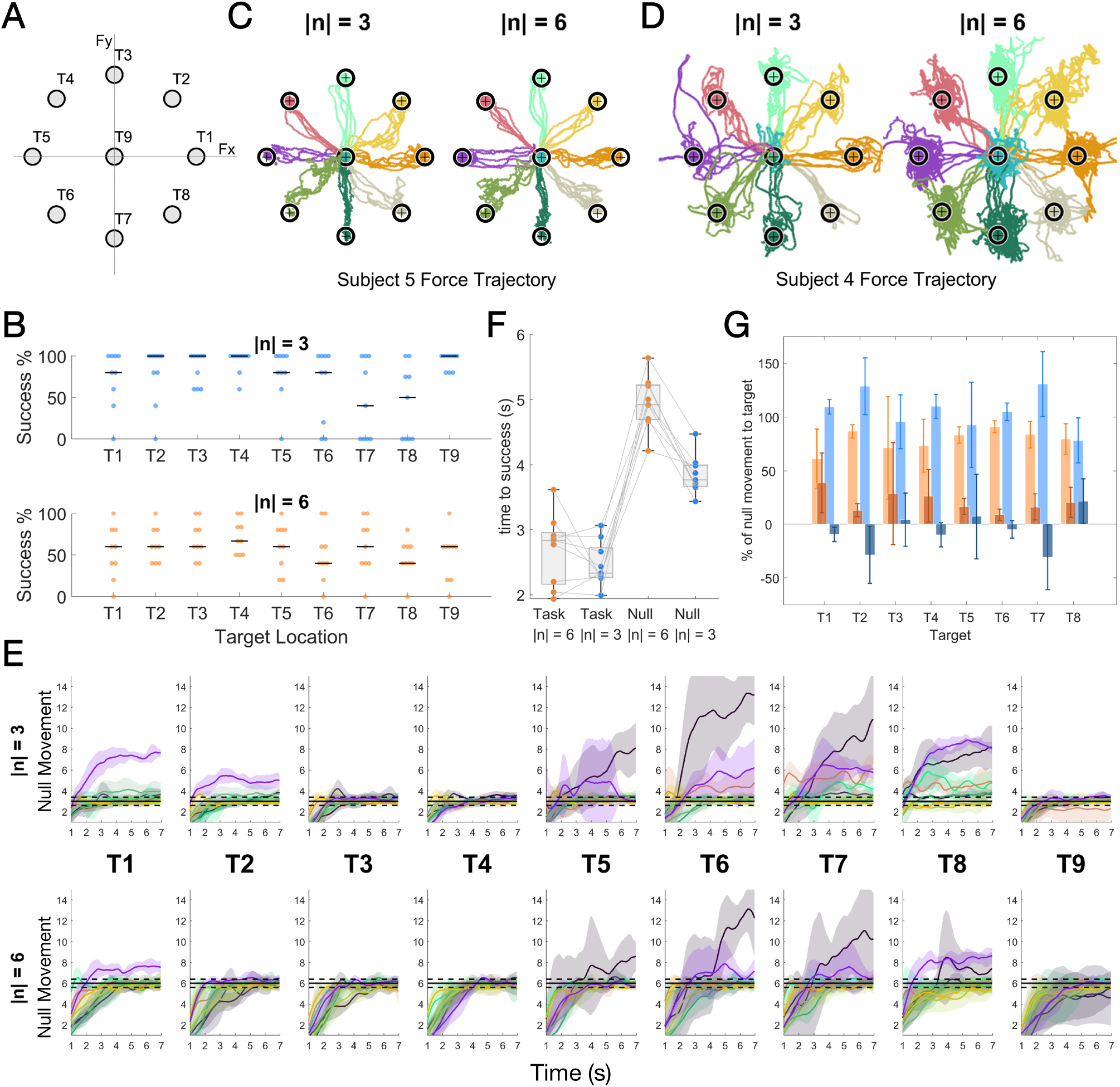
Performance during concurrent target reaching (Fig. 1F). (A) An illustration for the labels for targets in different locations. Note that for visual convenience the target size is not the same as was used in the session. (B) The success rate of different target locations and target sizes. The black horizontal lines indicate the median success rate. (C-D) Overlaid trajectories in the task-space for Subjects 5(C) and 4(D). Different colours depict the trajectories for different targets. (E) Amount of null-space movement in different target positions and sizes. Solid lines and the shades denote the mean and the standard deviation respectively of null-space movement by time. Black horizontal solid lines mark the target size and the black horizontal dashed lines denote the tolerance range for null-space movement. Each colour denotes 1 participant. (F) Target reaching time in the task- and null-space for different target sizes. (G) Percentage of null-space movement (median ± median absolute deviation) during phase 1 (light colours) and phase 2 (dark colours) for small (blue) and large (brown) targets. Negative values depict a reduction of the magnitude of null-space movement which can be due to overshooting in the null-space during phase 1.

Trajectories from the cursor position show that all target positions could be reached regardless of the target position and size, suggesting that there were no difficulties for the participants to move in the task-space (Fig. 4C-D). However, while some participants showed a relatively stable trajectory towards targets regardless of the target size (Fig. 4C), other participants showed much more variance in the task-space when higher co-contraction was required (Fig. 4D). This could be due to the different level of activity that different participants generated in the null-space while reaching for targets in the task-space. Null-space activity generated during target reaching for all participants suggest different reasons for failing to reach the targets. When the target was small, participants mainly failed the task due to generating too much null movement (T5-T8). For the larger sized targets, unsuccessful trials were caused by either insufficient null movement (T9) or also too much of the null movement (T6, T7) (Fig. 4E).

The latencies to successfully reach the target in task- and null-space evaluation clearly show that the target position was typically reached before the correct target size was obtained using null-space activity (Fig. 4F). The latencies depended on space and target size (Bayesian repeated measures ANOVA, *BF*_*inclusion*_ > 100 for space, target size and interaction of size and space) but there was no evidence for a difference between large and small targets for the task-space (post-hoc Bayesian paired t-tests with adjustment for multiple comparison; posterior odds>20 for all pairwise comparison except between ∥ **ñ**∥ = 3 and ∥ **ñ** ∥ = 6 for the task-space: posterior odds= 0.25; uncorrected *BF*_10_ = 0.6).

To investigate whether target reaching in the null- and task-space took place sequentially or concurrently, we split the null-space activity into 2 phases: Phase 1 lasted from movement onset until when the target in task-space was reached while Phase 2 lasted from when the target was reached in the task-space until when the target reach was completed (i.e. the target was reached in both task- and null-space). We used the EMG reading of the same target positions as in T1-8 taken during the calibration phase for each individual participant and predicted how much null-space movement they would have generated in reaching for the target in task-space without any additional co-contraction. This null-space movement was used as the baseline null-space movement, and the additional required null-space movement to reach the target size was then considered as the total null-space movement needed. From here we computed the proportion of null-space movement generated by the 2 phases as the contribution of the null-space movement to reach the target for the corresponding phase. For both target sizes, the contribution from Phase 1 was higher than from Phase 2 (Bayesian repeated measures ANOVA, *BF*_*inclusion*_ = 10, 0.7, 1.2 for phase, size and interaction),suggesting that target reaching mainly took place concurrently in the null- and task-space, and that the participants then fine-tuned the target size after reaching the correct target location (Fig. 4G).

## Discussion

We tested the capability of users to exploit the redundancy naturally present within the wrist’s musculoskeletal system as a source of signals for DoF augmentation. For successful DoF augmentation it is required that users can reliably control additional DoFs with minimal interference on their existing natural motion behaviours. In 2 sessions, we therefore evaluated if users could i) alter their wrist muscle activation patterns to reach for targets positioned in the task-intrinsic muscular null-space; and ii) simultaneously vary their activity in the null-space while performing isometric force reaching tasks. Our results show limited ability for participants to reliably control their behaviour within the muscular null-space as well as no clear improvement in the range of their null-space activation as a result of training. In contrast, while their performance was direction and target size dependent, the results show some potential for concurrent target reaching in the task- and null-space when standard co-contraction magnitude is used as a null-space input.

### Participants struggled to reach different targets in the null-space

Despite participants showing a range of null-space angles that they could manoeuvre the cursor to, only 2 out of 10 participants (Subjects 5 and 6) were able to progress beyond the third target in the NSTR task. Here because of the chosen target size, the fourth target would be the first target for which an initial strategy of moving to the first target’s centre location (as in a natural co-contraction) would be unsuccessful. The poor performance of most participants shows that they could not adapt their muscle activation patterns for reliable null-space target reaching within the given number and length of trials. This inability to reliably reach for targets in the null-space could have been caused by: i) variations in the participants’ natural co-contraction behaviour making it that the first target itself was difficult to reach; ii) participants not having enough repetitions and/or time to adapt their behaviour; and/or iii) the participants possessing insufficient flexibility and/or control of their muscle activation patterns.

4 out of 10 participants did not progress beyond the first target. This suggests that they were not able to reliably reproduce the natural co-contraction behaviour that they executed in the calibration phase. This might reflect that the natural co-contraction behaviour of each single participant possessed a sufficiently large variation such that while they might have been able to reach the target associated to their natural co-contraction, they could not hold the necessary muscle activation for the required 1 s. However, this is unlikely as some participants (for example Subjects 1, 3 and 9) displayed a consistent bias in their null-space activation within the target reaching phase (Fig. 2D) and never produced motion in the same direction as their original mean natural co-contraction activity. This bias may have been the result of an unobserved small postural change that occurred within the 2 minute exploration phase^26^, alternatively it could result from the change of the visualisation and the associated instruction when going from producing ‘natural’ co-contraction to change the cursor size during the calibration to the target reaching phase for which co-contraction now resulted in 2-dimensional motion. In either case, the participants did not show an ability to compensate for the presence of a bias nor for the natural variation in their wrist co-contraction.

The similar angle ranges (Fig. 3A) in the exploration and target reaching phases suggest that participants did not gain additional capability to reach angles in the null-space throughout the 80 trials of the target reaching phase. This indicates that at least over the time-scale of the experiment that there was no learning effect in the range of null-space activation behaviours. While it has been previously observed that atypical muscle activation patterns can be observed only with larger time for adaptation^15^, it is worth noting that both the motion efficiency and number of corrections (Fig. 3B,D) suggest that participants did not make use of extensive online adaptation when they were successful. This was despite the 7 s of reaching time being large enough that in some cases participants could return to the origin and make a second reaching attempt within the allotted time period.

Was there any difference that explained the better performance of the 2 most successful participants? There was no clear feature (for example experience with the device and/or demographics) that separated the 2 most successful participants from the other participants. However, with only 2 participants further data would be required to determine if there is a particular cause for these improved results, where for example these participants may simply possess a greater flexibility and natural control over their muscle activation patterns.

### Participants could simultaneously vary their co-contraction with task-space reaching

In the CTR session, we tested whether participants could vary the magnitude of their co-contraction in the null-space while simultaneously producing task-space force. The relatively high success rates shown in Fig. 4B and an even higher performance when consider only target reaching without holding (see Supplementary Figure S5) suggests that participants were capable of performing the task. However, their success rate was target position dependent. Among all the target positions, the success rate was lowest when the targets required force ulnar deviation. When isometric wrist movement in task-space was performed, it was also found to produce unintended natural null-space activity due to non-negativity of muscle activity. This natural null-space activity was found to be nonlinear across the 2D space and participant dependent. The null-space trajectories for different target positions in Fig. 4E suggests that the unintended natural null-space was larger in the ulnar direction (T6-8), causing the lower success rate in these directions. A more sophisticated nonlinear calibration that accounts for this natural null-space activity may be able to minimise this interference and enable greater success across these target locations.

The success rates were also target size dependent. Here for the small target size, failure was always the result of participants overshooting the target in the null-space, while for the large target size failure occurred due to both overshooting and undershooting the null-space magnitude. As a result of these 2 modes of failure, the median success rate was lower for the large target size (Fig. 4E). This indicates that the range and the accuracy of control of possible null-space activation levels during task-space motion is functionally constrained.

Despite taking longer to reach the targets in the null-space than in the task-space, the motion concurrency analysis (Fig. 4F,G) showed that most of the gross motion in the task- and null-spaces occurred simultaneously. The additional time to reach the null-space target therefore suggests that participants possessed less fine-motor control in their null-space than in their task-space actuation. Here, it is worth noting that participants have a lifetime of experience in performing task-space motions with fine-motor control. In contrast while natural co-contraction is used in natural movement (e.g., to “stiffen up” in response to disturbances^27^), the control of co-contraction within natural movement may take place less often and require lower accuracy than that of task-oriented force control. This may be a factor in the observed performance differences.

### Application considerations

Our null-space reaching results suggest that while users possess variability in their null-space behaviour, most lack the flexibility and control necessary for using all of the mechanical degrees of redundancy present in the wrist complex. This apparent inability to exploit novel muscle activation patterns differs from findings considering the entire upper limb^15^, where it has been observed that with sufficient time to make online motion corrections participants could learn to exploit atypical arm-wide muscle activation patterns for end-point reaching tasks. This suggests that while there is some potential for the use of musculoskeletal redundancy across an entire limb as a source of commands for movement augmentation it does not appear to be suitable at the level of the wrist joint, at least not within a short practice period on a single day. As a result, the number of possible signals that are directly available through the muscular null-space may be lower then previously thought.

It is however worth noting that while the wrist system was chosen as it is simple to measure in an isometric setup, due to difficulties in obtaining a reliable sEMG recording without cross-talk, the study only used 4 out of 5 wrist muscles after the exclusion of the extensor carpi radialis brevis (ECRB). Furthermore while no evidence of adaptation was found, the study only considered 1 session. The lack of an ECRB measurement limits the possible redundancy present within the wrist-complex such that there is only a 2-dimensional null-space, while the use of a single session limited the potential for possible motor learning. These 2 factors restricted the study’s ability to observe variation in muscle activation patterns, such that evaluation with all muscles (potentially through intra-muscular recording) or the consideration of other joints (such as the shoulder or the hip) that possess higher degrees of redundancy, are worth investigating over a multi-day experiment.

Our concurrent reaching results, in addition to those previously found with the upper limb^16^, indicate that variation of the magnitude of co-contraction may represent a mechanism for non-invasive augmentation control that can be manipulated in parallel with natural movement. However, in both our results and those of^16^, while participants could in general reach the different desired null-space magnitudes, the ability to hold that level of muscle activation was not as consistent and observably participant dependent. Furthermore, it is worth noting that because of a linear model assumption, as well as the non-negativity property of muscle activity, task oriented motion will always lead to activities in the 2D null-space. Here, the magnitude of co-contraction is also known to vary during natural motion for information exchange purposes^28^, such that natural co-contraction has a function and cannot necessarily be considered as a mechanism for augmentation by extension without there being new different muscle activation patterns observed.

Despite these concerns, the use of the muscular null-space is likely less application dependent than approaches that have considered the kinematic null-space^29^, for which the task-relevant kinematics changes for each different task. Since the primary cause of failure was the presence of involuntary null-space activation, the use of a participant dependent non-linear null-space that adapts the required co-contraction levels based on this involuntary activation could offer a mechanism of reducing any negative interference with natural motion behaviour while increasing individual participant success rate.

## Conclusion

Our results do not find evidence for a general ability to vary the wrist’s null-space muscle activation patterns. Furthermore we do observe that the wrist’s null-space activation patterns are coupled to task-oriented motions. Together these factors suggest limited potential for the use of the redundancy present within the wrist’s musculoskeletal system as a non-invasive source of signals for DoF augmentation.

## Acknowledgements

This research was supported in part by the European Commission grant H2020 NIMA (FETOPEN 899626) and Italian Ministry of Health grant (GR-2019-12370271).

