## Supplementary Material for "Control limitations in the null-space of the wrist muscle system"

[1]Bernstein Center Freiburg, University of Freiburg, 79104 Freiburg im Breisgau, Germany
[2]Faculty of Biology, University of Freiburg, 79104 Freiburg im Breisgau, Germany
[3]Department of Bioengineering, Imperial College of Science, Technology and Medicine, W12 0BZ London, UK
[4]Mechanical Engineering Department, the University of Melbourne, Victoria, Australia
[5]Center for Information and Neural Networks (CiNet), Advanced ICT Research Institute, National Institute of Information and Communications Technology, 1-4, Yamadaoka, Suita City, Osaka, Japan
[6]Laboratory of Neuromotor Physiology, IRCCS Fondazione Santa Lucia, Rome, Italy
[7]Department of Systems Medicine and Centre of Space Bio-medicine, University of Rome Tor Vergata, Italy
[8]Department of Biomedical and Dental Sciences and Morphofunctional Imaging, University of Messina, Italy
[*]MJL, JE should be considered joint first author
[**]CM, EB should be considered joint senior author

### Expanded calibration data

Each participant performed a calibration procedure consisting of *force/EMG calibration* and *null-space calibration* across both experimental sessions. Within these sessions, the participant specific *pulling vector matrix* and *'natural' co-contraction* directions were computed. The values for these variables are presented below:

#### Pulling vector matrix

Fig. S1 depicts the projection of the participants' pulling vector matrix into the flexion-extension and radial-ulnar deviation force, where each individual colour is associated with the effect of a different muscle recorded using surface EMG. It can be observed that there was variability in the identified participant pulling vectors between participants. However, the projections for an individual muscle typically occupied the particular region associated with the muscle's function (for example the extensors produce extension and the ulnaris muslces produce ulnar deviation). The regions that the muscle pulling vectors occupied appear to be consistent across the two sessions, with the exception of the FCR which appears to have changed between the NSTR and CTR sessions.

The quality of the fit of the linear mapping from surface EMG activity to measured force was evaluated through the *coefficient of determination* ($R^2$) resulting in $R^2 = 0.83 \pm 0.08$ and $0.87 \pm 0.05$ for the null-space target reaching and concurrent target reaching sessions, respectively. This suggests that while the linear approximation is accounting for much of the variability in the data, there is still some non-linearity not accounted for in the EMG-to-force mapping.

#### Natural co-contraction

Fig. S2 shows the average EMG recordings for each participant across the two different experimental sessions. Across subjects, our data showed little to no evidence for a deviation in their co-contraction EMG activities across the 2 sessions ($BF_{10} = 2.3, 1.6, 1.2, 0.5$ for ECRL, ECU, FCR, FCU, Bayesian Wilcoxon signed-rank test).

### Null-space target reaching failure quantification

As an additional investigation of the causes of failure during the null-space target reaching task, we studied a series of related metrics.

#### Minimum held distance and time in target

The proximity of a single trial to a successful reach was quantified through two metrics: the *minimum held distance* to the target and the *maximum held period* within the target. Here, the distance to the target was computed as the Euclidean distance between the participant's cursor position and the target location. The minimum held distance was then given by the smallest

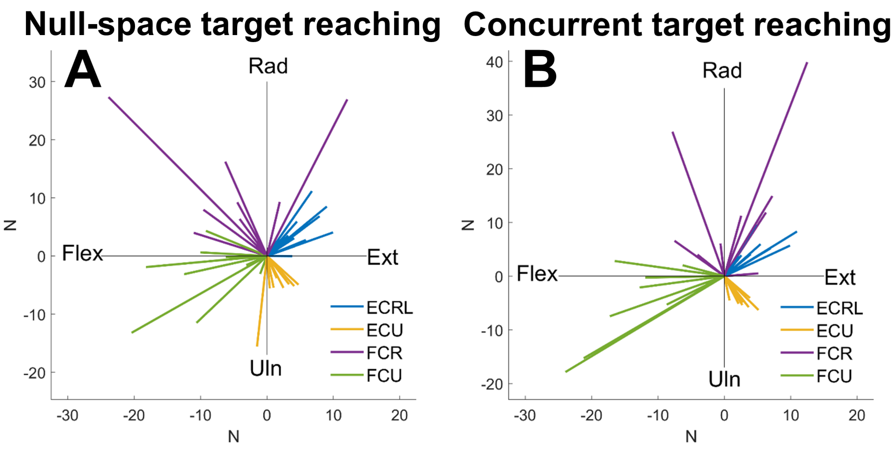


**Figure S1.** Pulling vector matrix visualisation. The flexion-extension and radial-ulnar deviation force produced by recorded activation of the flexor carpi radialis (FCR), flexor carpi ulnaris (FCU), extensor carpi radialis longus (ECRL) and extensor carpi ulnaris (ECU) muscles is visualised for the calibration conducted before the null-space target reaching session (A) and concurrent target reaching session (B). Each muscle is shown with a different colour, while individual pulling vectors are shown for each participant

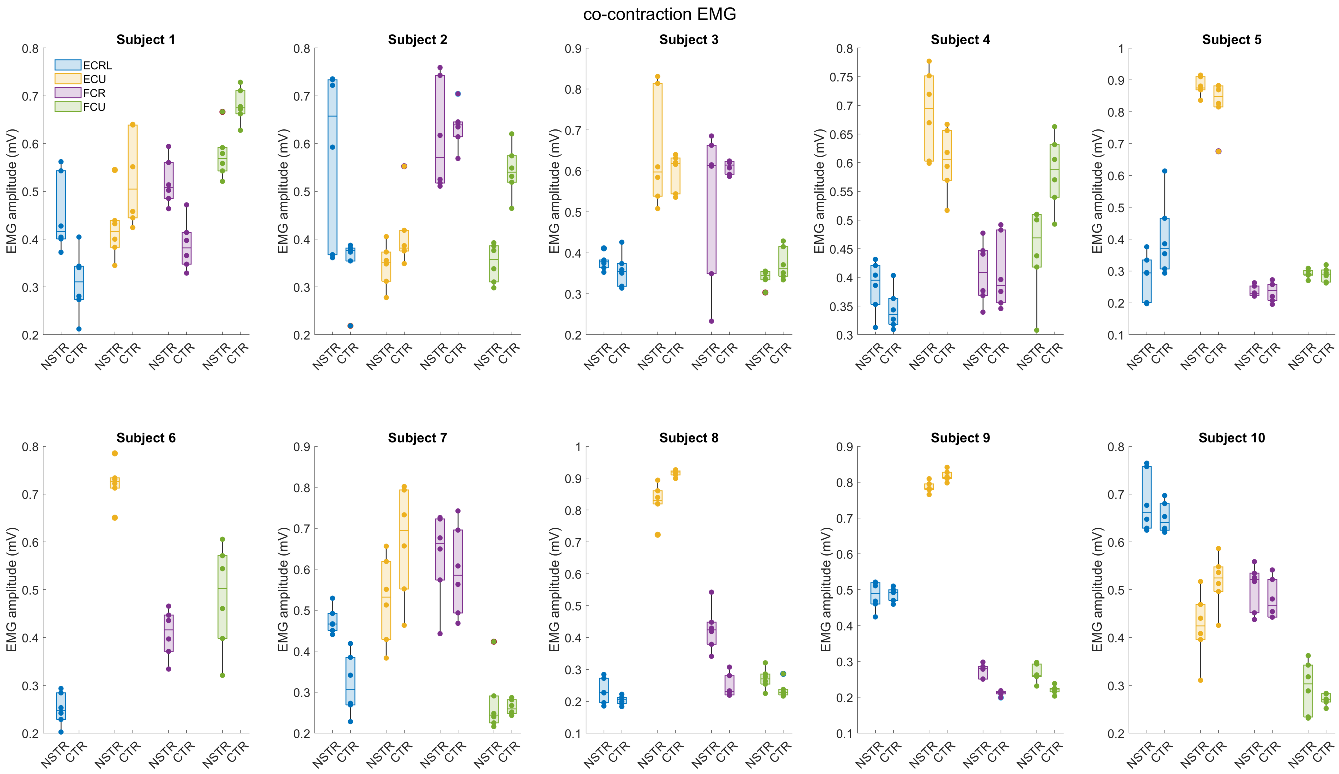


**Figure S2.** EMG activity during null-space calibration. For each participant, the average EMG activity for each muscle over the last 2 s of natural co-contraction is shown as a box plot with the single trial EMG activities also shown as individual dots. Two box plots are shown for each muscle corresponding to the recordings from the null-space calibration during the null-space target reaching (NSTR) and concurrent target reaching (CTR) sessions, respectively. Each muscle is shown in a different colour consistent with that of Fig. S1.

distance to the target that was held for 1 s without violating the force constraints. Fig. S3A shows the minimum held distance for each participant across the trials of the null-space target reaching session, where if there was no 1 s interval without force constraint violation no data was recorded. There did not appear to be a trend of participants reducing their minimum held distance before successful trial reaches.

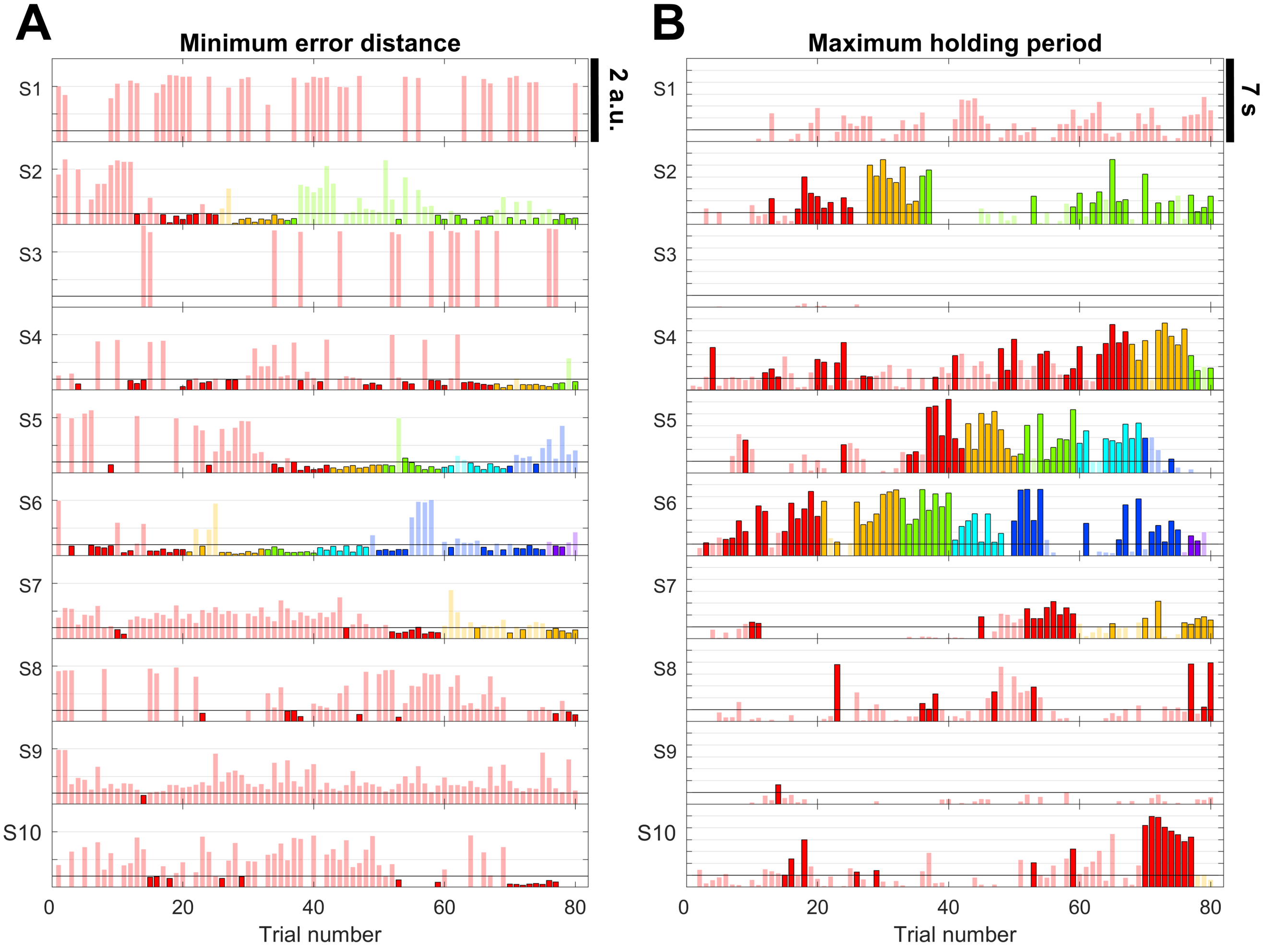


**Figure S3.** Minimum held distance and time in target for each trial. (A) The minimum held distance (without violating the force constraint) that was held over a 1 s time window consistent with the required holding condition. Here, the black horizontal line depicts the required distance for a successful trial. (B) The maximum held period over the trial for which the participant had the cursor within the required distance threshold from the target without consideration of the force constraint. Here, the black horizontal line depicts the required holding time for trial success. Saturated colours depict successful trials and transparent colours illustrate unsuccessful trials. Each colour is coded for an individual target. Colour representation is the same as in Figure 2 and 3E in the main text.

The *maximum held period* (Fig. S3B) was instead computed as the maximum period of time in which the participant kept the cursor within the threshold distance of the target location consistently without interruption, where if a cursor was never in the target then no data was recorded. Failure of a trial could be due to unsuccessful target reaching / holding from a cursor or violation of the force constraints. From these results it can be observed that participants were able to in 53.4% of the failure trials (318 trials from the total number of trials across all subjects) reach the target and in 22.5 % of the failure trials (134 trials from all subjects) they could also hold their null-space position to be within the target for more than the required 1 s holding time. However, that this was often coupled to force production such that the individual trial was not successful.

### Effect of threshold conditions

From Fig. S3B, it was observed that there were a number of trials for which the participants were able to reach the target but not hold their position. Fig. S4A further illustrates the participant trial-by-trial success at reaching without holding. It can be seen that if the success condition did not include holding, then Subjects 2, 7, 8 and 9 would have all at least reached one further target location (due to having 80% success out of a window of ten trials). However, Subjects 1 and 3 would still not progress beyond the first target location.

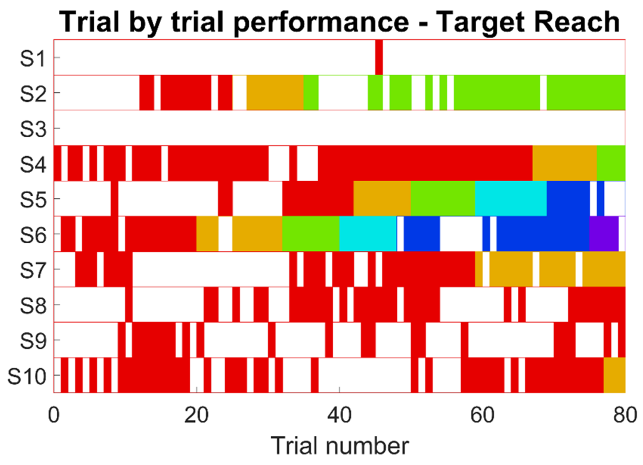


**Figure S4.** Trial failure visualisation. Distribution of trials where the participant reached (without necessarily successfully holding) the target location for each participant. A reached trial is shown by a coloured bar, where each colour represents a different target location. Without the holding requirement, 8 out of 10 participants would have met the criteria to move past the first target.

### Additional concurrent target reaching performance metrics

#### Reaching versus holding performance

Fig.S5A shows the participants success rate for the concurrent reaching if the 1 s holding condition was not considered. From this figure it can be seen that participants could typically reach all targets in both the task- and null-space regardless of the size or location. Most failures in the experiment were therefore reflective of an inability to hold the null-space position rather than an inability to reach it.

#### Null-space histogram

Fig. S5B depicts the range of null-space activation (as a histogram) that the participants produced for each task-space target location. For each individual participant, the direction of the null-space activation appears to remain within a consistent range across the different target locations and size. This indicates that participants used a similar strategy of EMG activation that was independent of the task-space requirements as well as the required physical exertion.

### Force variability

We investigated the impact of the wrist-joint muscle redundancy control on subsequent task-space reaching by evaluating the difference between the force/EMG calibration 20% scaled maximum voluntary force target reaching that participants performed before (pre) and after (training) for each experiment session. We then analysed the force variability during the force hold phases and compared it between session (pre or post) and target for each experiment (Fig. S6). For the null-space target reaching task we found strong evidence against there being a change of force variability (as measured by the median absolute deviation) with respect to the target or the interaction between the target and session (Bayesian repeated measures ANOVA, $BF_{inclusion} = 0.006, 0.001$ respectively) and moderate evidence against any change across the pre and post sessions ($BF_{inclusion} = 0.324$). Similarly, for the concurrent target reaching we found strong evidence against there being a change of force variability with respect to the target or the interaction between the target and session (Bayesian repeated measures ANOVA, $BF_{inclusion} = 0.037, 0.006$ respectively) and moderate evidence against any change across the pre and post sessions ($BF_{inclusion} = 0.236$).

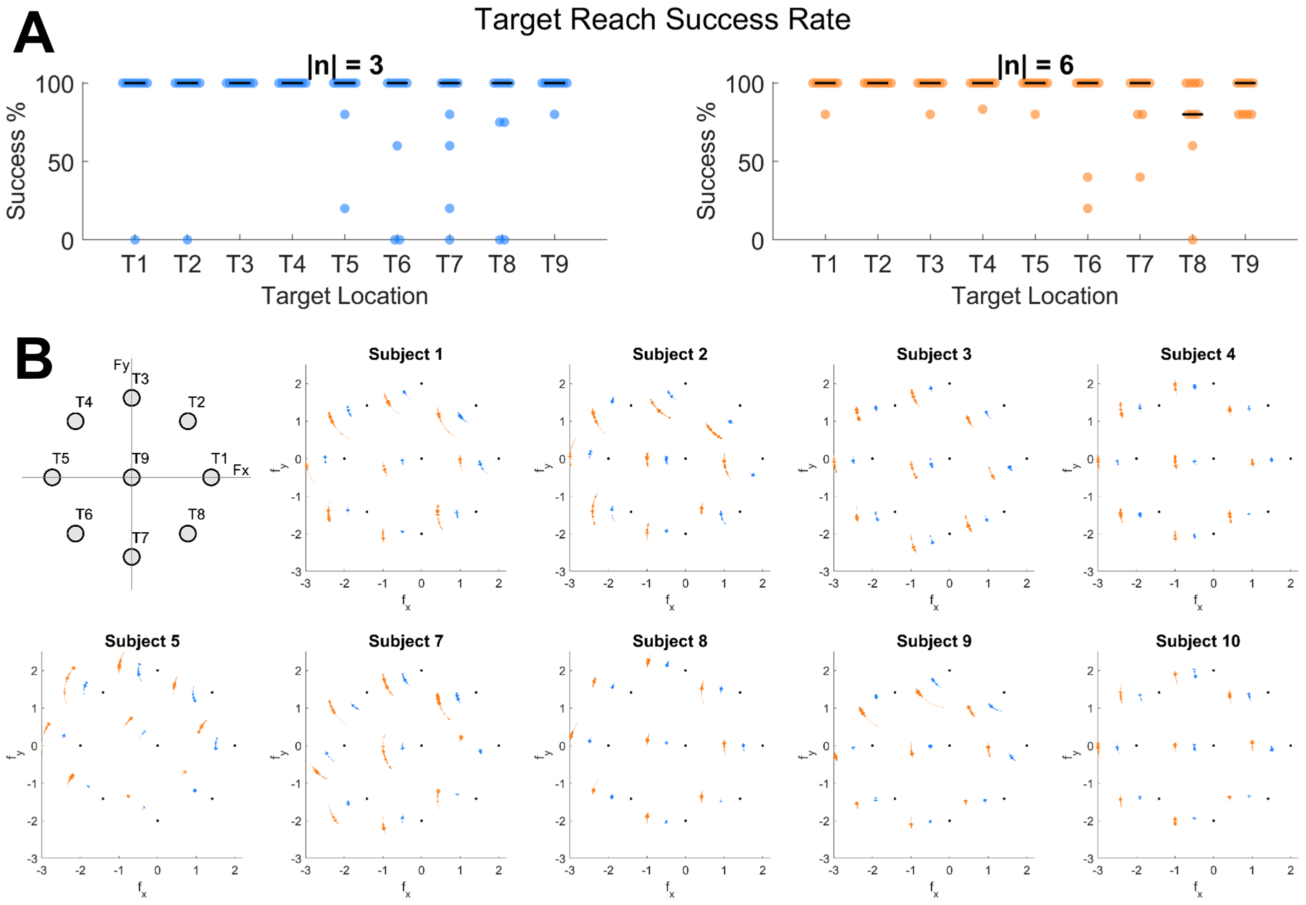


**Figure S5.** Additional concurrent target reaching performance results. (A) The reaching rate of different target locations and target sizes. (B) Histogram showing the angles reached for each target location. Angular coordinate shows the reached angles for each target where blue is used for target sizes $\|\tilde{\mathbf{n}}\| = 3$ and orange is used when $\|\tilde{\mathbf{n}}\| = 6$. Width shows the frequency that the angles were reached. Note that due to a computer crash, Subject 6's data was not saved during the CTR session and is therefore not included.

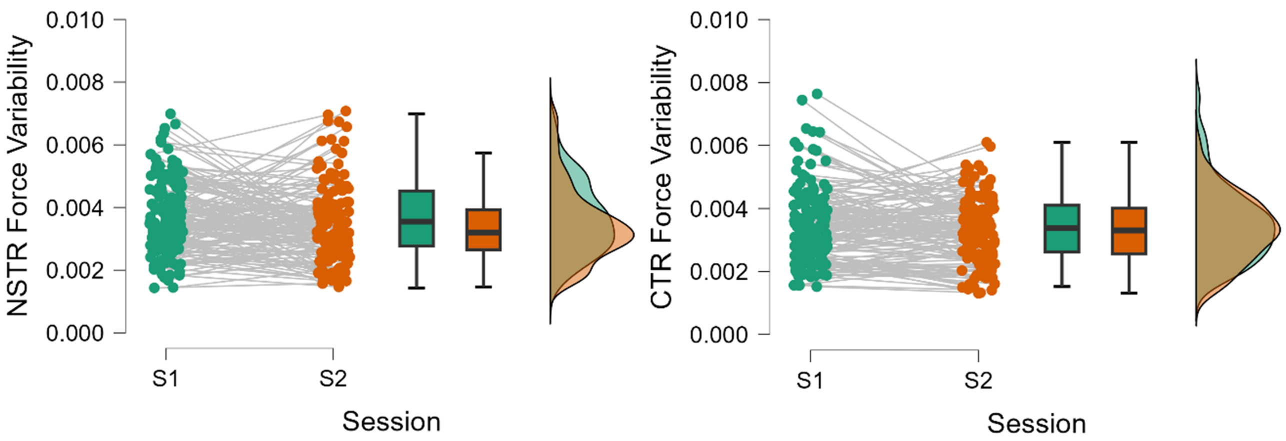


**Figure S6.** Force variability during task-space isometric target force production (within the force/EMG calibration) at 20% MVC. Force variability was quantified by the median absolute deviation of the force during the hold period and was assessed directly before (pre) and after (post) both experiment sessions (NSTR on the left, CTR on the right). Dots depict each individual target from each participant.
